# Influence of plant defense signaling and innate insect metabolic differences to the overall performance of fall armyworm (*Spodoptera frugiperda*) corn and rice strains on maize as a host

**DOI:** 10.1101/2023.12.07.570551

**Authors:** Bhawana Israni, Sabine Haenniger, Bettina Raguschke, Michael Reichelt, Jonathan Gershenzon, Daniel Giddings Vassão

**Author notes:** (BI); (DGV) Max Planck Institute for Chemical Ecology, Jena, Germany.

## Abstract

The fall armyworm (FAW, *Spodoptera frugiperda*) is a well-known crop pest that feeds mainly on grasses. Separate strains are known to infest maize (corn) and rice that show varying degrees of developmental and metabolic differences, as well as reproductive isolation. Here we show that the greater performance of the corn compared to the rice strain on maize leaves may be explained by several factors. Maize plants respond to herbivory by the rice strain with greater levels of defense hormone signaling and greater accumulation of defensive benzoxazinoids. Moreover, measurements of the activity of a glucosyltransferase involved in benzoxazinoid metabolism and the transcript levels of the encoding gene revealed that the corn strain had higher benzoxazinoid detoxification potential than the rice strain. The two strains also exhibit constitutive differences in the expression of an alternate variant, with potential consequences for differential regulation of the glucosylation activity. These factors may account for the better performance of corn strain larvae on maize leaves, perhaps in combination with the other differences we found in maize defense metabolites after FAW herbivory by untargeted metabolomics.

## Introduction

The fall armyworm (FAW, *Spodoptera frugiperda* J.E. Smith, Lepidoptera: Noctuidae) is a highly cosmopolitan insect pest that has spread from North and South America throughout the world owing to its highly migratory nature. In accordance with its extensive spread, this pest has been documented on more than 350 plant species, although it is best known to feed on agriculturally significant grasses such as maize (corn), rice and sorghum. Interestingly, a marked level of genetic differentiation has been reported between FAW populations isolated from corn and rice fields (Kergoat et al. 2012; Oliveira et al. 2022), and these have accordingly been named as the corn strain and the rice strain. Corn strain larvae are often associated with corn, sorghum and cotton plants, while the rice strain association is more prominent in fields of rice and forage grasses. The strains display differences in development (Groot et al. 2010; Meagher Jr et al. 2004; Meagher and Nagoshi 2012), differences in metabolic activity (Hay-Roe, Meagher,Nagoshi 2011; Veenstra, Pashley,Ottea 1995) and mating barriers in nature (Groot et al. 2010), facilitating host-associated specialization. Although the strains show clear differentiation based on mitochondrial and sex-linked genetic markers, some exceptional instances of mixing between corn and rice strains in the field have also been reported (Schlum et al. 2021).

The ability of insect herbivores to specialize on a host plant may play a significant role in speciation and adaptive radiation (Nylin, Slove, Janz 2014). Yet we often know little about the biochemical and physiological mechanisms underlying specialization. For the corn and rice FAW strains, it has been suggested that salivary effectors can differentially induce the host response towards herbivory, leading to the observed differences in herbivore performance (Acevedo et al. 2018). Insect saliva, regurgitant, frass or eggs, often mediate the very first interactions with the host (Mattiacci, Dicke, Posthumus 1995; Ray et al. 2016; Tian et al. 2012). These products then trigger the biosynthesis of defense hormones, notably jasmonic acid (JA), salicylic acid (SA), ethylene (ET) and abscisic acid (ABA), enabling the plant to respond characteristically to a given herbivore. The JA pathway leads to the biosynthesis of the JA-isoleucine conjugate (JA-Ile), the prime ligand for the CORONATINE-INSENSITIVE1 (COI1)-jasmonate ZIM-Domain (JAZ) receptor complex, activating COI1 dependent WRKYs and other transcription factors that subsequently leads to synthesis of plant defense metabolites (Schweizer et al. 2013). Differences in plant signaling pathways can thus modulate defense metabolites in different ways and thus shape the performance of different herbivores.

A large variety of plant defense metabolites can be induced by hormone signaling pathways in response to herbivory in maize, including terpenoids, phenolics, and a variety of amino acid-derived compounds. Among the latter are the benzoxazinoids (BXDs), tryptophan-derived defenses found in many grasses that are toxic to a wide range of chewing, piercing-sucking and root herbivores (Wouters et al. 2016). Stored as glucosides, BXDs are activated by glucosidase action on plant damage. The ability of different FAW strains to specialize on maize could depend on the extent to which they induce BXDs on attack, but more information is needed on this question. Furthermore, the capability of FAW strains to feed on maize could depend on their resistance to BXDs. In this view, we have shown that FAW can detoxify the active BXD aglucones, such as 2,4-dihydroxy-7-methoxy-1,4-benzoxazin-3-one (DIMBOA), by stereospecific re-glucosylation catalyzed by a by a UDP-<u>g</u>lycosyltransferase (UGT) designated SfUGT33F28, resulting in a stable, biologically inert product (Israni et al. 2020; Wouters et al. 2014). Corn strain larvae have been shown to express constitutively higher levels of *SfUGT33F28* transcripts than rice strain larvae, which may lead to greater amounts of re-glucosylated DIMBOA in their frass. But more work needs to be done to understand strain differences in detoxifying BXDs. Recently, we have demonstrated that *SfUGT33F28* can be expressed as transcriptional variants that are regulated by DIMBOA and can interact with the canonical protein and regulate the basal activity of SfUGT33F28 (Israni et al. 2022). It would be thus interesting to learn how alternate *SfUGT33F28* transcripts differ between FAW strains.

In this work, we try to understand which factors might cause FAW strain performance differences on maize leaves by comparing defense signaling and BXD accumulation in the plant in response to herbivory by the corn vs. rice strain. In addition, we search for other metabolic changes and determine inter-strain differences in detoxification.

## Results

### Rice strain herbivory leads to greater jasmonate signaling than corn strain herbivory

To obtain detailed insights into the differences in maize phytohormone signaling in response to the two FAW strains, targeted measurements were carried out to measure jasmonates in maize plants subject to different treatments: mechanical wounding and herbivory by either the corn or rice strain of the FAW. Unwounded control maize plants were kept as controls.

FAW herbivory clearly led to higher accumulation of JA and its active isoleucine conjugate JA-Ile after herbivory as compared to mechanically-wounded and unwounded control plants (Figure 1). Within the first hour of herbivory, JA and JA-Ile showed an average induction of 2.18- and 3.02-fold respectively compared to unwounded controls (Figure 1A, 1B). However, strain-specific differences could not be ascertained. Differences between the corn and rice strains could be observed after prolonged caterpillar feeding (24 h) such that rice strain herbivory led to a stronger induction of the jasmonate pathway overall in maize leaves compared to corn strain herbivory. Prolonged rice strain herbivory led to an average 3.74- and 4.97-fold induction of JA and JA-Ile respectively compared to unwounded control (Figure 1A, 1B). Corn strain herbivory, on the other hand, led to a 2.18- and 3.86-fold induction of JA and JA-Ile respectively compared to unwounded control (Figure 1A, 1B).

**Figure 1.**
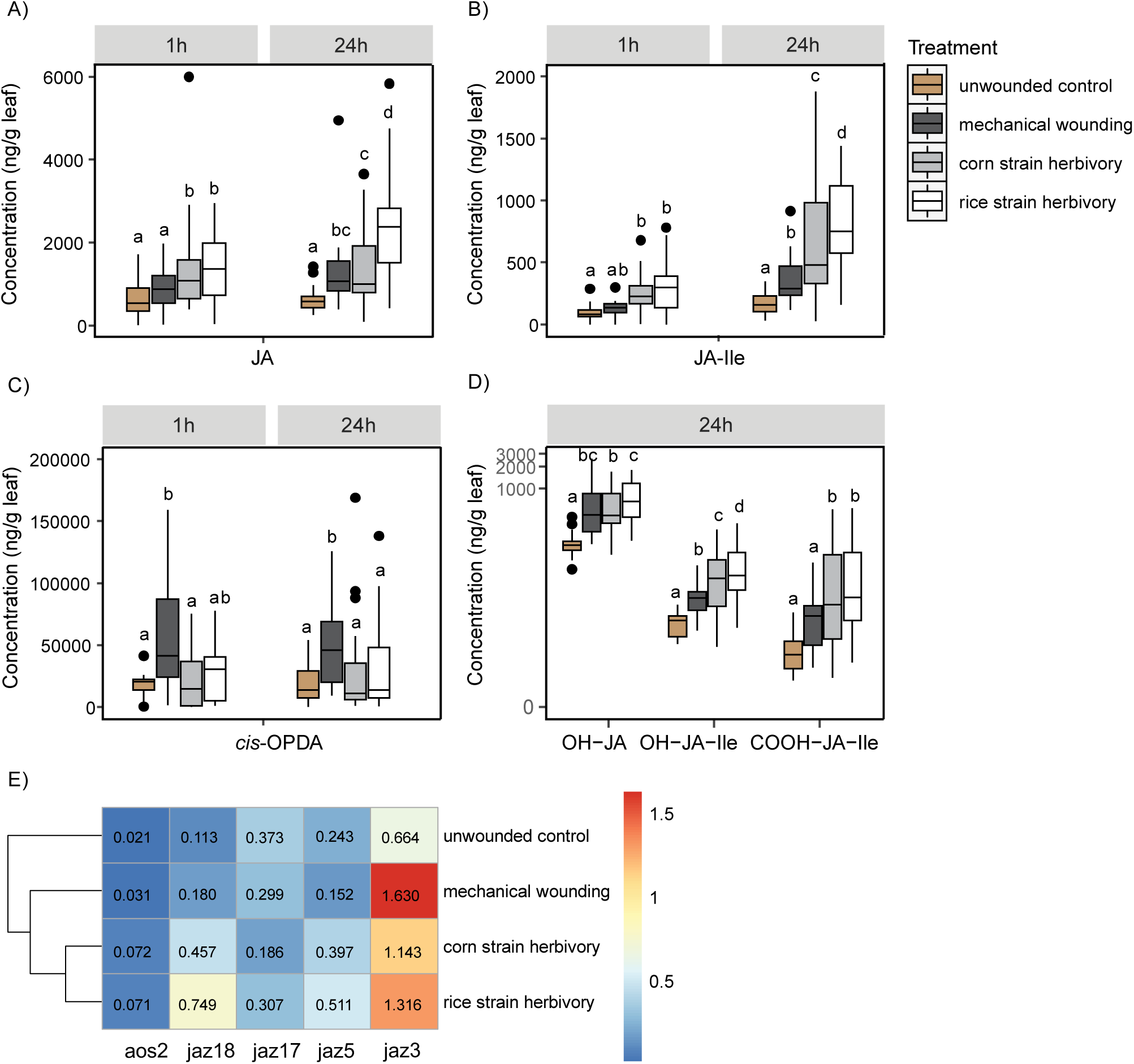
Differential JA signaling induced in response to FAW corn and rice strain herbivory. Accumulation of A) JA; B) JA-Ile conjugate; and C) *cis*-OPDA after 1 h and 24 h of herbivory (n = 30-40). D) Accumulation of JA metabolites 24 h post herbivory (n = 30-40). E) Heatmap showing differential expression of maize genes in response to herbivory. Maize actin was used as a housekeeping gene. Rows were clustered using the Pearson method (n= 20-25). Two-way ANOVA was performed and Holm Sidak method was applied to perform pairwise comparisons (A-D). All data were square root (A-C) transformed/ log transformed (D) to meet the criteria for normality for conducting statistical comparisons. Small letters on the plots represent significant differences (P< 0.05). JA, jasmonic acid; JA-Ile, jasmonic acid isoleucine conjugate; *cis*-OPDA, *cis*-12-oxophytodienoic acid; OH-JA, 12-hydroxyjasmonic acid; OH-JA-Ile, 11/12-hydroxyjasmonic acid isoleucine conjugate; COOH-JA-Ile, 11/12-carboxyjasmonic acid isoleucine conjugate.

Since further metabolism of JA has also been shown to be regulated in a stress-specific manner (Fernández-Milmanda et al. 2020), we measured several other JA metabolites 24h after herbivory began, and found that these mimicked the trend observed with the previously measured compounds (Figure 1D). Accumulation of hydroxylated JA was noted to be 4.4-fold upon rice strain herbivory compared to the unwounded control, significantly higher than that observed for corn strain herbivory (3.3-fold compared to unwounded control plants). Hydroxylated JA-Ile was found to be 5.1-fold higher in plants subject to corn strain herbivory and 6.7-fold higher in plants subject to rice strain herbivory compared to unwounded control plants. Accumulation of carboxylated JA-Ile was noted to be 13-fold higher upon corn strain herbivory when compared to the unwounded control and 15.5-fold higher upon rice strain herbivory compared to the unwounded control, although statistically significant strain differences were not present. Taken together, these measurements indicate a somewhat greater response of JA defense signaling from the rice strain than the corn strain when feeding on maize plants.

The differences observed in jasmonate accumulation between the treatments were also reflected in their transcriptional responses to herbivory (Figure 1E). The increases in jasmonates in plants subject to FAW herbivory were accompanied by an elevated expression of the gene encoding the biosynthetic enzyme allene oxide synthase (*Zm00001d028282*, *AOS2*) with increase upon herbivory of an average of 2.3-fold compared to mechanically-wounded plants and an average of 3.5-fold compared to unwounded control plants. This result was in accordance with earlier reports on FAW herbivory on maize (Acevedo et al. 2018). We also analyzed the transcripts of *JAZ* genes, which encode proteins that act as transcriptional repressors of the JA pathway, focusing on those found to be highly expressed in the maize leaves and most inducible with MeJA, 24 hours post treatment (Sun et al. 2021). *JAZ18* (*Zm00001d020614*, ZIM28/TIFY25; named after the conserved domains) was significantly induced in plants subject to corn and rice strain herbivory when compared to the untreated control plants (average 4-and 6.65-fold) or mechanically wounded plants (average 2.55- and 4.2-fold. *JAZ17* (*Zm00001d020409*, ZIM1/TIFY24) expression was lower in maize plants subject to mechanical wounding or herbivory compared to the unwounded controls (0.5-to 0.8-fold), with the most down-regulation seen in plants subject to corn strain herbivory. *JAZ5* (*Zm00001d033050*, ZIM18/TIFY9) was weakly up-regulated in response to corn and rice strain herbivory (1.6-fold and 2.1-fold respectively), but non-significantly when compared to controls. Both *JAZ18* and *JAZ5* transcripts were earlier reported to be induced in response to the plant elicitor peptide Zmpep3 (Huffaker et al. 2013), but *JAZ5* was shown to respond more generally to both Zmpep3 as well as *N*-linolenoyl-l-glutamine, a fatty acid-amino acid conjugate elicitor reported from oral secretions of many lepidopterans. *JAZ3* (*Zm00001d029448*, ZIM24/ TIFY10B) was up-regulated 1.7-to 2-fold in response to herbivory and 2.45-fold in response to wounding alone. Overall, maize plants subject to FAW herbivory cluster separately compared to mechanically wounded and unwounded control plants with stronger responses in general after rice strain herbivory when compared to corn strain herbivory.

### Herbivory by rice and corn strains induces differential expression of benzoxazinoid pathway genes and benzoxazinoid accumulation

Targeted analyses were carried out to determine the effects of FAW herbivory on the formation and accumulation of benzoxazinoid (BXD) defenses in maize, since BXD aglucones have been previously shown to impact FAW performance (Israni et al. 2020). A scheme for the maize BXD biosynthetic pathway is depicted in Figure 2A. Quantities of the most abundant BXD aglucones in maize, 2,4-dihydroxy-7-methoxy-1,4-benzoxazin-3-one (DIMBOA), 2,4-dihydroxy-7,8-dimethoxy-1,4-benzoxazin-3-one (DIM_2_BOA), 6-methoxy-2-benzoxazolinone (MBOA) and 6,7-dimethoxy-2-benzoxazolinone (M_2_BOA) exhibited significant differences within the first hour of herbivory.

**Figure 2.**
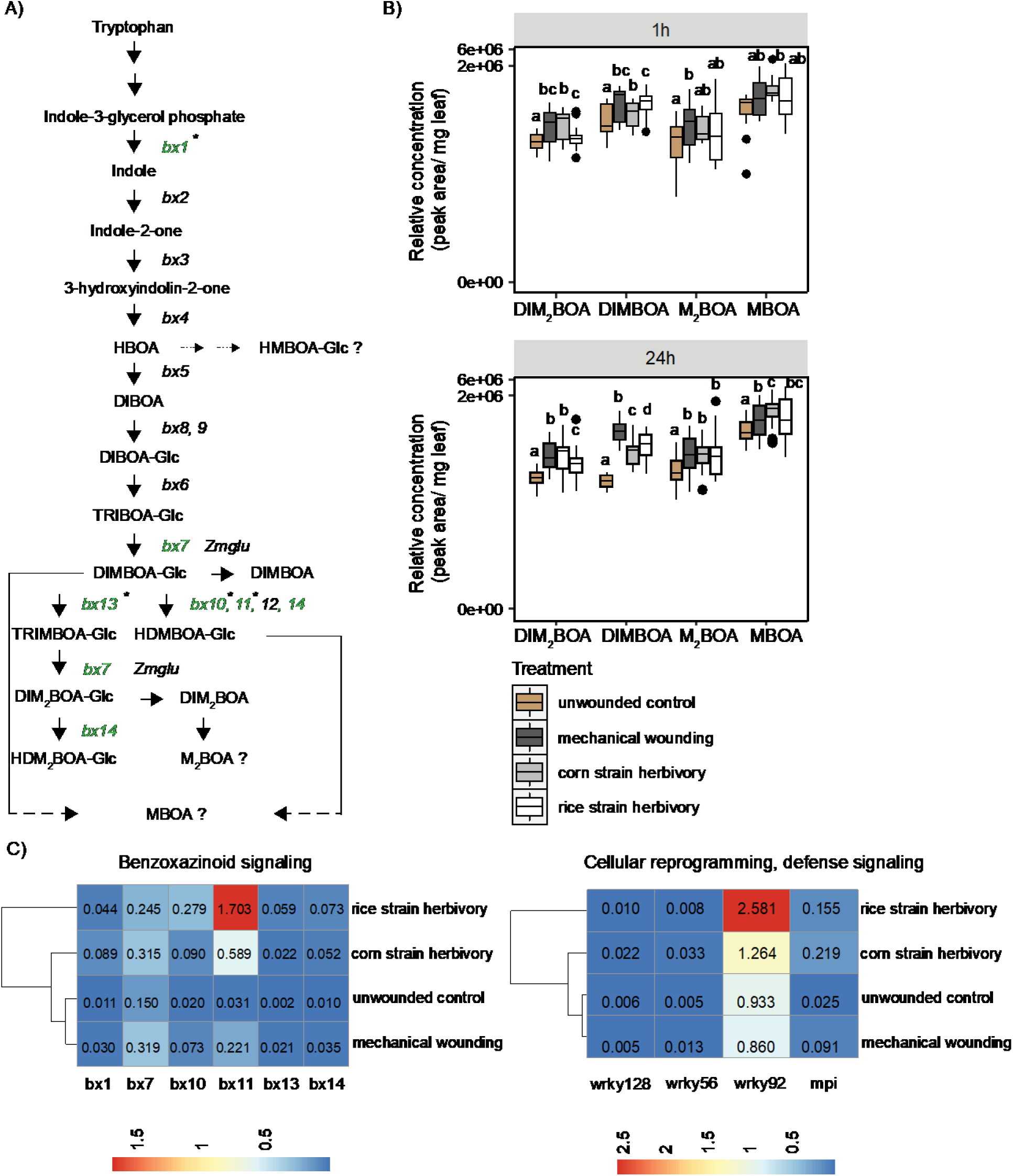
Accumulation of benzoxazinoids (BXDs) and upregulation of BXD biosynthetic genes in maize plants in response to FAW herbivory. A) Schematic showing BXD biosynthetic pathway with genes upregulated in response to FAW herbivory highlighted in green (* indicates significant differences, P< 0.05), B) BXD aglucone accumulation in maize leaves subject to FAW herbivory expressed as relative concentration (peak area/mg leaf) (n= 30-40), C) Heatmaps showing differential expression of maize BXD biosynthetic and other regulatory genes in response to herbivory (n= 20-25). Maize actin was used as a housekeeping gene. Rows were clustered using the Pearson method. Two-way ANOVA was performed and Holm Sidak method was applied to perform pairwise comparisons. All data were log transformed (A-B)/ square root transformed (C) to meet the criteria for normality to conduct statistical tests. Small letters on the plots represent significant differences (P< 0.05). DIMBOA, 2,4-dihydroxy-7-methoxy-1,4-benzoxazin-3-one; DIM_2_BOA, 2,4-dihydroxy-7,8-dimethoxy-1,4-benzoxazin-3-one; MBOA, 6-methoxy-2-benzoxazolinone and M_2_BOA, 6,7-dimethoxy-2-benzoxazolinone.

FAW herbivory generally led to significant increases in BXD accumulation compared to unwounded control plants. Within the first hour of herbivory, there was a 3.76-fold induction for DIM_2_BOAin response to corn strain herbivory and a 1.52-fold induction in response to rice strain herbivory; and DIMBOA, with 1.3-fold induction observed in response to corn strain and 2.67-fold induction in response to rice strain herbivory (Figure 2B, upper panel). An average of 3-to 4-fold induction was observed for accumulation of MBOA and 1.7- to 3.2-fold for M_2_BOA in leaves subject to FAW herbivory, but strain-specific differences were not observed (Figure 2B, upper panel). Over a 24-hour period of herbivory, further BXD accumulation could be observed, with DIM_2_BOA found to be on average 6.5-fold induced upon corn strain herbivory relative to unwounded control plants in comparison to an average 4-fold induction upon rice strain herbivory. DIMBOA accumulation was noted to be 9.26-fold upon corn strain feeding on maize leaves relative to unwounded control plants, compared to a 13.73-fold induction upon rice strain feeding (Figure 2B, bottom panel). Interestingly, upon mechanical wounding alone, DIMBOA induction upon mechanical wounding alone was more pronounced compared to FAW herbivory. Statistically significant differences were not detected for MBOA and M_2_BOA. A trend for differential M_2_BOA accumulation at 24 h, however, could be observed in line with that seen at 1h with a 2.71-fold higher accumulation upon corn strain herbivory and 4.92-fold higher accumulation upon rice strain herbivory in comparison to unwounded plants (Figure 2B, bottom panel). Patterns for the relative abundance of the benzoxazinoid glucosides in the maize leaves across different treatments are provided in **Supplementary figure 1**. Strain specific differences could be observed in only a few cases.

Subsequently, expression profiling of the BXD biosynthetic genes was carried out to understand the molecular response to herbivory by the two FAW strains (Figure 2C, left panel). From the core BXD pathway, *Bx1*, responsible for catalyzing the first committed step of BXD biosynthesis (Frey et al. 2009), was upregulated 8-fold upon feeding from corn strain caterpillars, but only 4-fold upon feeding by rice strain caterpillars compared to the unwounded control plants. *Bx7*, which catalyzes the conversion of TRIBOA-Glc to DIMBOA-Glc was up-regulated in response to herbivory as well as to mechanical wounding alone (1.6- to 2.1-fold), with no strain differences. For the latter part of the BXD biosynthetic pathway, we observed strong induction of *Bx10*-*14* transcripts. DIMBOA-Glc-O-methyltransferases *Bx10* and *Bx11*, responsible for conversion of DIMBOA-Glc to HDMBOA-Glc (Meihls et al. 2013), were also up-regulated in response to both mechanical wounding and FAW herbivory, with up-regulation of 4.4-fold in response to corn strain herbivory compared to unwounded controls, and 13.3-fold in response to rice strain herbivory. *Bx11* was the most up-regulated gene overall in response to rice strain herbivory, and was highly induced when compared to unwounded control (∼ 50-fold induction) and mechanically wounded plants (∼ 7.7-fold). *Bx13*, a 2-oxoglutarate-dependent dioxygenase, which catalyzes the conversion of DIMBOA-Glc to TRIMBOA-Glc (Handrick et al. 2016) was also up-regulated in response to herbivory, with a highly significant induction observed for rice strain-treated plants (∼ 34—fold compared to unwounded control), while mechanical wounding alone led to a 12.4-fold induction of *Bx13* transcripts. TRIMBOA-Glc formed as a result of *Bx13* activity can further be O-methylated by *Bx7* activity to form DIM_2_BOA-Glc, and both DIM_2_BOA-Glc and DIMBOA-Glc in turn can serve as substrates for the O-methyltransferase *Bx14* leading to generation of HDM_2_BOA-Glc and HDMBOA-Glc which are known to be induced in maize upon herbivory (Handrick et al. 2016). Consistent with these results, we also observed elevated *Bx14* levels upon mechanical wounding (3.43-fold) and herbivory-with a 5.14-fold induction observed for corn strain-treated maize plants when compared to unwounded controls and a 7.2-fold induction noted for rice strain-treated plants. A heatmap showing all the gene expression data for different treatment groups is shown in Figure 2C. The genes contributing to the strain differences are highlighted in the BXD biosynthetic scheme shown in Figure 2A. Again, maize plants subject to rice strain herbivory clustered distinctly compared to all other treatments.

Differences in gene expression were also observed for maize homologs of transcription factors previously implicated in triggering host defenses against herbivory (Palmer et al. 2019; Tzin et al. 2017), leading to an overall transcriptional reprogramming that could induce various defensive compounds in maize leaves after FAW feeding (Figure 2C, right panel). The putative switchgrass (*Panicum virgatum* L.) homolog of *AtWRKY72* was previously reported to be up-regulated in response to FAW feeding in two types of cultivars tested (Palmer et al. 2019). Furthermore, *Zm00001d039532* (*ZmWRKY56* in maize GDB, *Arabidopsis* ortholog *AtWRKY72*) and *Zm00001d010399* (*ZmWRKY92* in maize GDB, *Arabidopsis* ortholog *AtWRKY33*) were recently reported to be up-regulated in response to Zmpep3 treatment along with a host of other *ZmWRKY*s that were induced (Poretsky et al. 2021). In our experiments, *Zm00001d017444* (*ZmWRKY128* in maize GDB, *Arabidopsis* ortholog *AtWRKY12*) and *Zm00001d039532* were significantly up-regulated in the leaves subject to corn strain feeding, but only weakly induced in leaves subject to rice strain feeding. *ZmWRKY128* was up-regulated 3.35-fold in plants subject to corn strain herbivory compared to untreated control maize plants, while *ZmWRKY56* was up-regulated ∼6.9-fold upon corn strain herbivory. *ZmWRKY92*, on the other hand, was induced in the leaves upon corn strain herbivory (∼1.35-fold with respect to controls) compared to rice strain herbivory (∼2.8-fold with respect to controls) (Figure 2C). The *mpi* gene, on the other hand, encodes a maize proteinase inhibitor that accumulates in response to wounding and herbivory (Tamayo et al. 2000) and is known to inhibit the insect’s digestive enzyme repertoire. In line with previous reports, the induction of *mpi* transcripts was higher in plants subject to herbivory (8.9-fold in leaves subject to corn strain treatment with respect to the unwounded control and 6.32-fold in leaves subject to rice strain treatment) than mechanical wounding alone (3.71-fold), although statistically significant strain-specific differences were not ascertained.

Altogether, maize plants subject to rice strain herbivory clearly cluster separately compared to corn strain herbivory as well as mechanically wounded and unwounded control plants in reference to the expression of transcription factors involved in plant anti-herbivore defense.

### Untargeted metabolomics reveals induction of several maize metabolites in response to herbivory by the corn and rice strains

To determine additional changes in the maize leaf metabolism that occur in response to FAW herbivory, we analyzed leaf extracts with an untargeted approach using liquid chromatography– quadrupole time of flight-mass spectrometry (LC-qToF-MS). Analyses were carried out to compare damage by FAW corn and rice strains with mechanically-wounded and unwounded plants after 1 and 24 hours. More than 600 features were detected in the negative mode. Multivariate data analysis using orthogonal partial least squares discriminant analysis (OPLS–DA) was performed to highlight the differences among the groups and metabolites putatively involved (Figure 3). The validity of the OPLS-DA models thus generated was confirmed by performing permutation analysis (n= 100) to minimize the risk of over-fitting.

**Figure 3.**
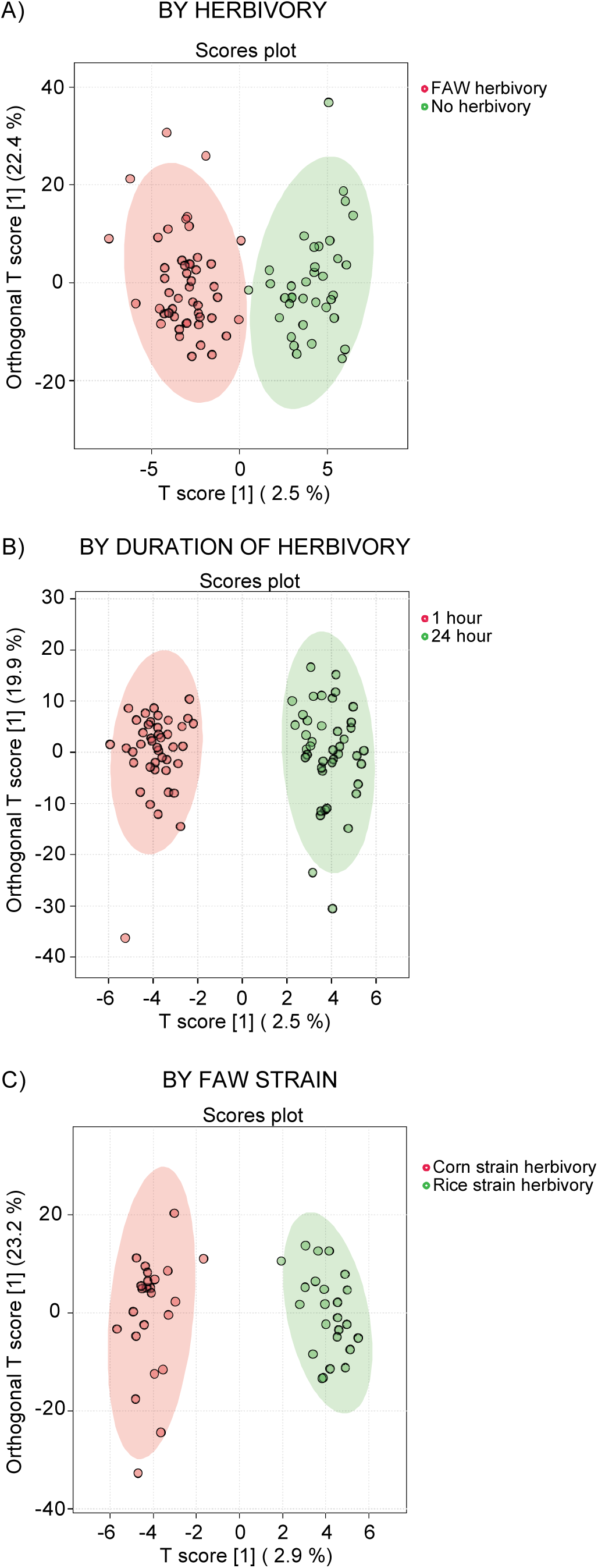
Orthogonal partial least squares discriminant analysis (OPLS–DA) of variation in maize leaf metabolome in response to various FAW herbivory treatments (n=9-12). On the OPLS-DA scatter plot, the horizontal axis explains variation between the groups tested, while the vertical axis (orthogonal score) explains variation within the groups. All analyses were performed using Metaboanalyst 5.0. The data were subject to log normalization, and pareto-scaling (mean-centered and divided by the square root of the standard deviation of each variable).

An overview of the major changes in individual metabolites together with the SIRIUS annotation (Dührkop et al. 2019) is presented in Table 1. In addition to the anticipated changes in the accumulation of JA and other jasmonates, several fatty acid derivatives were found to be enhanced in response to herbivory, with the most significant induction observed for hydroxylated fatty acids 18:3 -2OH and 18:3 -3OH in the leaves subject to rice strain herbivory. Several phenolics, amino acids and amino acid conjugates, were also found to be differentially accumulated in response to rice or corn strain herbivory or when compared to control plants. Notable induction was observed for a coumaroyl tyramine conjugate and caffeoyl 3-hydroxy tyrosine in maize leaves subject to corn strain herbivory compared to rice strain herbivory. Further work is need to establish the chemical identity of these features and to ascertain their role in response to FAW herbivory.

**Table 1.**
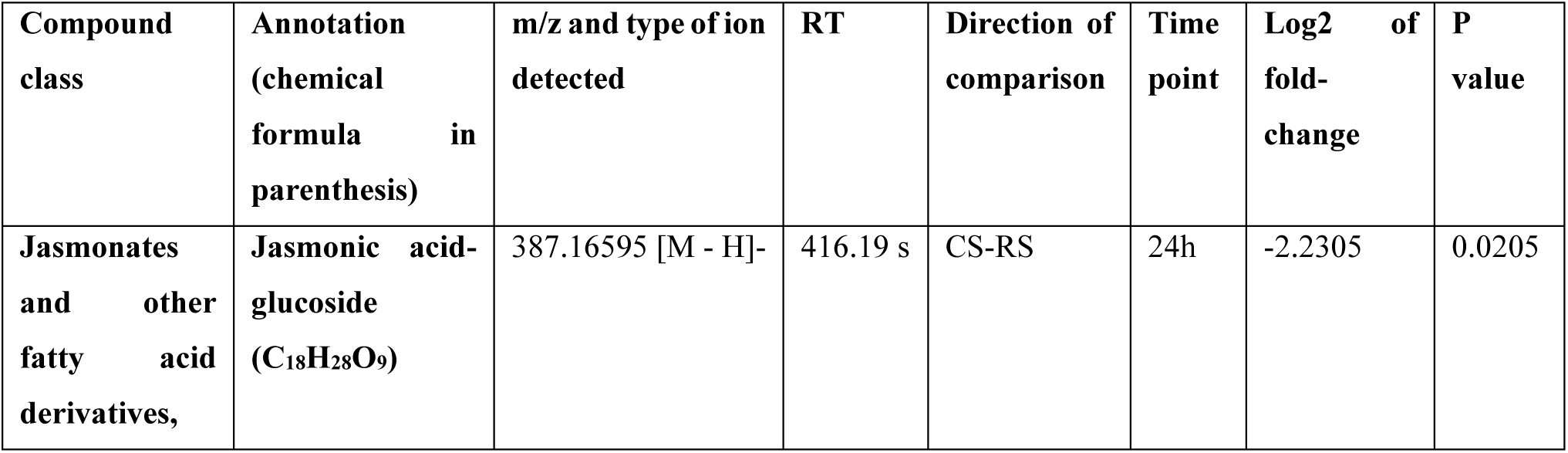
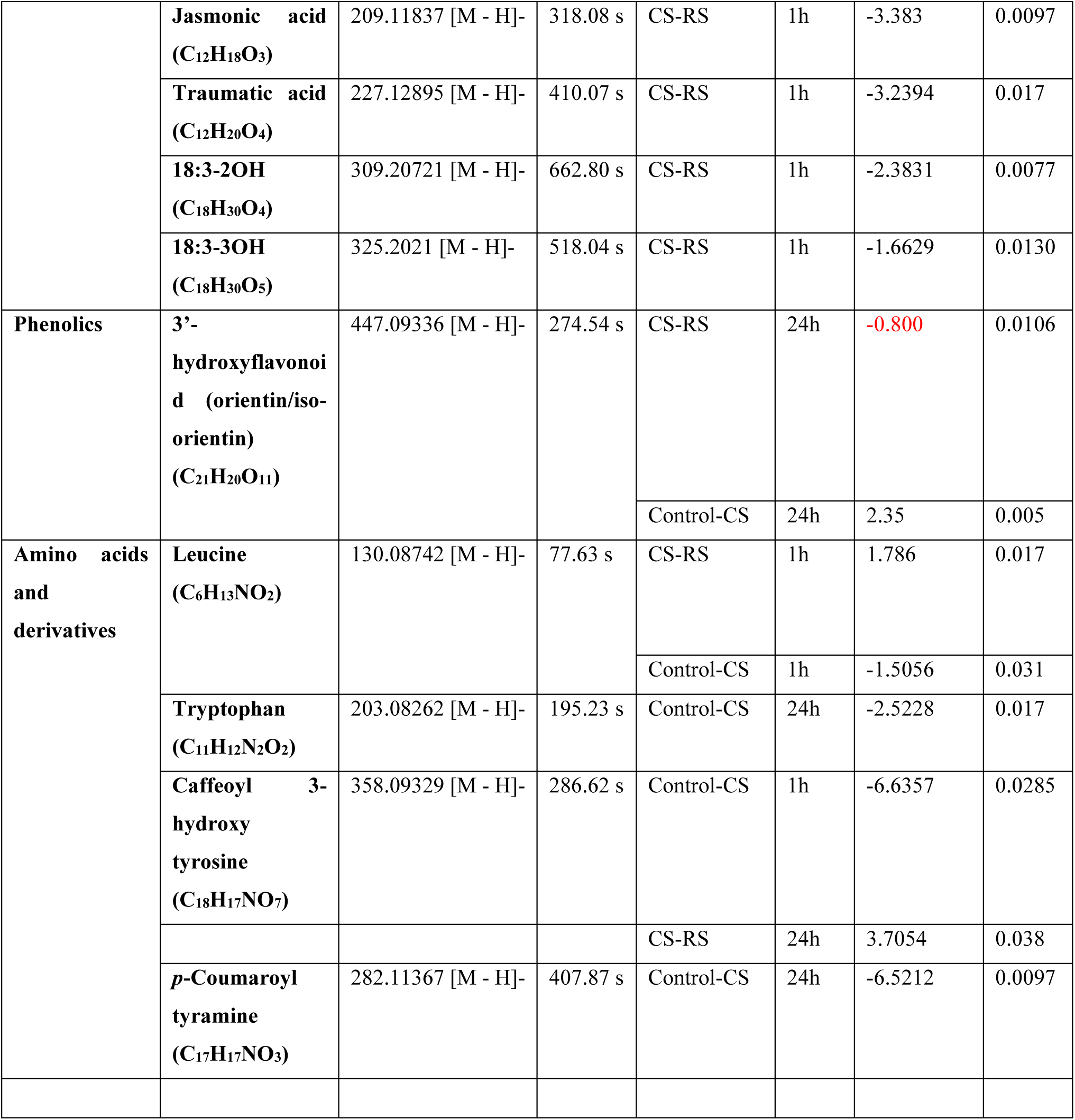
Most significant metabolomic features that were altered in maize leaves subject to herbivory by the two FAW strains. Fold-change (FC) refers to the differences in the qToF intensity of the features and was calculated based on a volcano plot with a fold-change threshold of 2 using Metaboanalyst 5.0 (fold change highlighted in red does not meet the cut off set for analysis). Annotations of compound identity are from SIRIUS. CS, corn strain; RS, rice strain.

### FAW strains display variability in benzoxazinoid detoxification activity and transcriptional regulation of the underlying detoxification gene

Another factor that could be responsible for differences between the ability of corn and rice strains to feed on maize leaves is their differential ability to detoxify BXD defense compounds. Since there is already considerable information on the ability of FAW UDP-glucosyltransferases (UGTs) to detoxify BXDs by stereoselective re-glucosylation, we investigated the potential of corn and rice UGTs to detoxify DIMBOA, one of the principal BXDs in maize.

We fed caterpillars from two populations of each of the two FAW strains on either a pinto bean-based artificial diet or on maize leaves. Corn strain larvae clearly showed higher enzymatic capability for glucosylation towards the BXD aglucone DIMBOA when compared to rice strain larvae, in line with previous observations (Israni et al. 2020) (Figure 4A, left panel). The trend among the individual populations of each strain analyzed is depicted in Figure 4A (right panel). Corn strain population CS-1 showed a clearly high activity towards the aglucone when compared to another corn strain population CS-2 (2.21-fold higher), and the other rice strain populations (2.75- to 7.7-fold higher). Among the rice strain populations, RS-1 displayed a much higher activity for DIMBOA (2.8-fold) compared to RS-2, which ranked the lowest amongst all populations analyzed.

**Figure 4.**
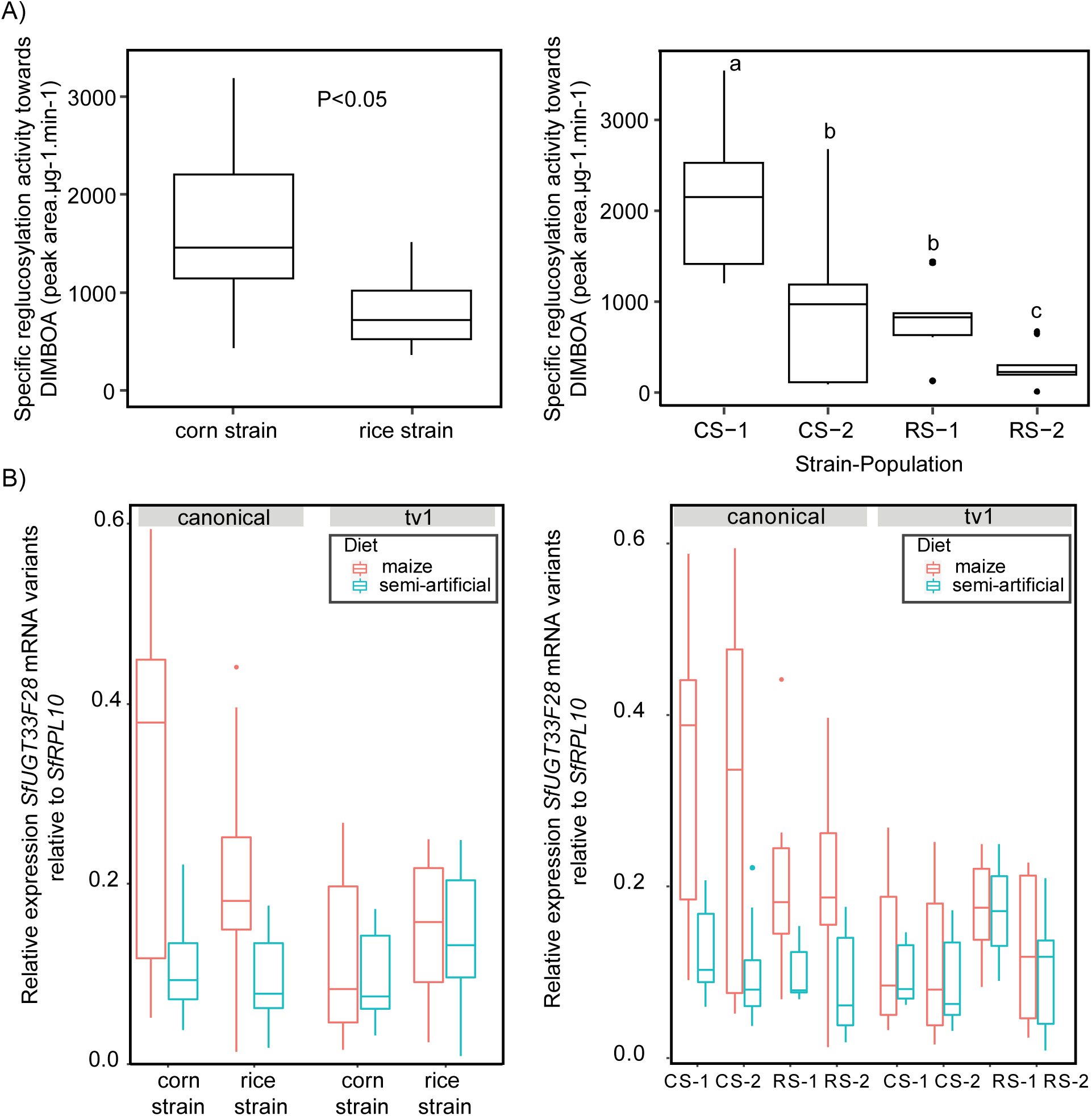
(A) Comparison of UDP glucosyltransferase activity from gut extracts of different strains tested towards DIMBOA with respect to strain (left panel) and population (right panel), B) Comparison of expression of canonical and variant (*tv1*) transcripts of the UDP glucosyltransferase gene *SfUGT33F28* with respect to strain (left panel) and population (right panel) tested on different diets (n=10). *SfRPL10* was used as a housekeeping gene. Two-way ANOVA was performed for A) and Holm Sidak method was applied to perform pairwise comparisons. Small letters on the plots represent significant differences (P< 0.05). The linear mixed effects model for B) was fitted using Restricted Maximum Likelihood (REML) approach. CS-1, corn strain population 1 (PRC); CS-2, corn strain population 2 (COB); RS-1, rice strain population 1 (FLR), RS-2, rice strain population 2 (ONA).

At the transcript level, we studied *SfUGT33F28* gene, which had been previously shown to encode a UGT that re-glucosylated BXDs in the FAW gut (Wouters et al. 2016; Wouters et al. 2014). Furthermore, an alternate SfUGT33F28 variant was discovered that is formed by alternative splicing and is inducible by DIMBOA. This variant is truncated, but interacts with full length SfUGT33F28 to enhance re-glucosylation (Israni et al. 2022). Accordingly, both the canonical *SfUGT33F28* and alternately spliced transcript *SfUGT33F28_tv1* were measured from the RNA isolated from insect guts and normalized to housekeeping gene *SfRPL10*. (Figure 4B). While the canonical transcripts were overall higher in the corn strain populations CS-1 and CS-2 than in the rice stain populations, the alternative transcripts were found to be constitutively higher in rice strain population RS-1 compared to all other populations studied. Higher levels of variant transcription in the RS-1 larvae might account for the higher activity observed towards DIMBOA compared to that in the other rice strain population. A mixed effects model was computed to understand these relationships, and the relative expression of *SfUGT33F28* was indeed found to be a function of transcript type (canonical or alternately spliced, Pr(>|t|)= 2.64e-10), diet (pinto bean-based or maize, Pr(>|t|)= 1.95e-10), strain (CS/ RS, Pr(>|t|)= 0.002606) as well as the population under consideration (intercept; Pr(>|t|)= 1.63e-09) (Figure 4B). The interactions between the variables transcript type and diet (Pr(>|t|)= 2.40e-05), transcript type and strain (Pr(>|t|)= 0.000318), diet and strain (Pr(>|t|)= 0.018516) contributed significantly to the model tested. These relationships can be summarized in the following model as: Relative *SfUGT33F28* expression∼ Transcript type x Diet x Strain + (1|Population) (***)

## Discussion

FAW is a highly cosmopolitan lepidopteran pest that attacks various economically important agricultural taxa, with a particular feeding preference for grass crops. The damage inflicted on Africa’s staple crop maize, one of the highly preferred host plants for FAW, is reported to contribute to yield losses in the range of 21-67% (Day et al. 2017; Kumela et al. 2019). Although the FAW originated in sub-Saharan Africa, it has colonized many other regions in recent years, including much of the grain growing belt of Asia. Two strains of FAW have been reported to occur in sympatry within the native range-the corn strain and rice strain, originally named after the fields from which these insects were sampled. Although the two strains are morphologically identical, they show development differences to varying degrees according to the host plant (Acevedo et al. 2018) and exhibit differences in their mating behavior (Groot et al. 2008). The strains exhibit various pre- and post-zygotic mating barriers, and the females of these strains display variable pheromone compositions (Cruz-Esteban et al. 2018; Groot et al. 2008). This information needs to be combined with knowledge of their induction of resistance to plant defenses to better understand the evolution and possible control of the FAW.

Comparing the two strains, the corn strain usually performs much better than the rice strain on maize, although a few reports show the opposite trend (Meagher, Nagoshi,Stuhl 2011). While both FAW strains have previously been shown to cause differential induction of gene transcripts pertinent to JA pathway in maize and accumulation of proteinase inhibitors (Acevedo et al. 2018), the rice strain generates a more pronounced transcriptional response in maize owing to differences in salivary phospholipase C activity, leading to much reduced larval growth. This suggests that strain-derived cues or behavior can modulate maize defenses.

Here we showed that rice strain herbivory indeed leads to an elevated defense response in maize compared to the corn strain owing to greater accumulation of JA and JA-Ile. Jasmonates have been demonstrated to be crucial to maize defense against insect attack, in particular against other *Spodoptera* species (Yan et al. 2012). An elevated defense response is evidenced by greater accumulation of jasmonates and an up-regulation of *ZmAOS2* transcripts. From the two clades of AOS that have been described-AOS1 and AOS2, it has been suggested that the AOS2 clade is involved in the production of JA directly due to its ability to catalyze the conversion of either 9-HPOD or 13-HPOT into the corresponding allene oxides (Borrego and Kolomiets 2016).

JA signaling can also be regulated through metabolism of JA-Ile via 12-hydroxylation and subsequent hydrolysis and silencing of genes pertinent to this reaction have been already shown to have consequences for resistance to FAW (Chung et al. 2022). In our study, the amounts of 12-hydroxy JA and 12-hydroxy JA-Ile were higher after 24 hours of rice versus corn strain herbivory. On another level, the JA signal transduction process is ultimately realized by the action of JA-Ile. As a ligand associated with the F-box protein COI1 (coronatine insensitive 1), JA-Ile mediates the binding of the JAZ (jasmonate-zim domain) repressor proteins to this complex. This leads to the targeted degradation of the JAZ proteins, freeing transcription factors that can now direct the expression of various JA-dependent genes, including those belonging to the MYC (master regulator of cell cycle entry) bHLH (basic helix loop helix), MYB (myeloblastosis viral oncogene homolog), EIN3 (Ethylene insensitive 3, regulator of ethylene signaling), and DELLA (regulator of gibberellic acid/GA signaling) families. For instance, induction of jasmonate signaling in maize triggers the degradation of MYB11 and ZML2 (TIFY family), leading to the de-repression of the lignin biosynthetic gene encoding caffeic acid O-methyl transferase (*Comt*) (Vélez-Bermúdez et al. 2015). *Comt* promoters of grasses were in general demonstrated to be enriched in AC-GAT(A/C) *cis*-regulatory elements to which TIFY family members can bind, suggesting a mode of regulation for these proteins in lignin pathway. The differences in JA signaling after herbivory by the FAW corn and rice strains are reflected in our own results showing a greater expression of JAZ18 in the rice strain, in accordance with a previous report on *ZmJAZ18* (designated as *JAZ1a* in this publication). (Han and Luthe 2022).

Following herbivory by the two FAW strains, we also observed variability in the expression of WRKY transcription factors, often involved in phytohormone signaling and response to stress. *AtWRKY12* from *Arabidopsis* was previously found to be involved in repression of lignification and secondary cell wall development (Wang et al. 2010), while its equivalent in maize *ZmWRKY128* was more upregulated by corn strain over rice strain herbivory, which might partially explain the better performance of corn strain larvae compared to rice strain larvae. The *Arabidopsis AtWRKY72*, on the other hand, has been shown to cause repression of *AOS* transcription by inducing DNA hypermethylation in its binding site in the *AOS* promoter and so attenuate JA synthesis (Hou et al. 2019). The very significant increase of ∼6.9 fold in the transcript of the corresponding maize WRKY(*ZmWRKY56*) subject to corn strain herbivory is in line with the relatively lower JA signaling in maize leaves subject to corn strain herbivory compared to rice strain herbivory. Finally, the *Arabidopsis AtWRKY33* has been shown to positively regulate JA- and ET-mediated defenses (Kundu and Vadassery 2021; Zheng et al. 2006) with a well-supported role in wounding and herbivory. An up-regulation of the maize homolog *ZmWRKY92* during rice strain herbivory over corn strain herbivory is in accordance with the increased JA defense response towards the rice strain.

BXDs make up another significant fraction of the maize leaf metabolome induced in response to FAW herbivory. The corresponding aglucones, which result from the activity of ꞵ-glucosidase during herbivory are detrimental to insect development. Accumulation of maize BXDs has already been reported to occur in response to JA pathway activation upon wounding, elicitation with the elicitor peptide, ZmPep (Huffaker, Dafoe, Schmelz 2011; Huffaker et al. 2013), and FAW herbivory in general (Glauser et al. 2011). The aglucones DIMBOA (greater after rice strain herbivory) and DIM_2_BOA (greater after corn strain herbivory) seem to contribute to most to significant strain-specific differences, while MBOA shows a trend for enhanced abundance in leaves subject to rice strain herbivory. Accordingly, *bx7*, responsible for the biosynthesis of DIMBOA-Glc is significantly up-regulated in response to FAW herbivory. JA is especially known to promote the conversion of DIMBOA-Glc into HDMBOA-Glc (Oikawa, Ishihara, Iwamura 2002; Oikawa et al. 2004). Accordingly, in our study the biosynthetic enzymes catalyzing this reaction, encoded by *Bx10*, *11* and *14*, were all highly induced in response to rice strain relative to corn strain herbivory.

Untargeted metabolite analysis further revealed that rice strain herbivory induces a higher accumulation of various fatty acid derivatives-such as 18:3 -2OH and 18:3 -3OH in the very first hour of herbivory than the corn strain. Accumulation of fatty acids and their derivatives is a general stress response in plants particularly after wounding and herbivory (Banerjee and Roychoudhury 2020; Kallenbach et al. 2011; Marti et al. 2013). Dodecenoic and dodecanoic acid derivatives are generated by the hydroperoxide lyase pathway utilizing linoleic (18:2D9,12; 18:2) and α-linolenic (18:3D9,12,15; 18:3) acids. Lipoxygenases (LOXs) generate 13S-hydroperoxy dienoic and trienoic acids from linoleic and linolenic acids that are in turn converted by hydroperoxide lyase to C_12_ aldehydes and the C_6_ green leaf volatiles. Green leaf volatiles such as hexanal and (3Z)-hexenal, and compounds such as(9Z)-traumatin produced as a result of these reactions can in turn undergo many other modifications, generating multiple defense signals, which may contribute to the greater defense reaction in response to rice strain herbivory (Grechkin 2002).

Other defensive compounds induced in maize in response to FAW herbivory included hydroxycinnamic acid derivatives including *p*-coumaroyltyramine and caffeoyl 3-hydroxytyrosine. Both these compounds were found to be highly induced in response to corn strain herbivory as compared to undamaged control plants. Both *p*-coumaroyltyramine and feruloyltyramine have been previously reported to accumulate in tomato, tobacco and maize plants following wounding (Ishihara et al. 2000; Pearce et al. 1998), in tobacco following salicylic acid treatment and fungal and viral pathogens (Sun et al. 2019), and in maize following *S. littoralis* herbivory (Marti et al. 2013). However, artificial diet experiments with *p*-coumaroyltyramine showed no effect on insect performance. *S. littoralis* larvae were in fact able to convert a majority of *p*-coumaroyltyramine to other metabolites that supported its growth (Marti et al. 2013). Similar utilization of *p*-coumaroyl tyramine by the FAW corn strain could explain its overall better performance on maize than the rice strain.

The performance differences between the two FAW strains may also be ascribed to differential ability to detoxify the different classes of metabolites induced in maize upon herbivory. This aspect was investigated in more detail utilizing the well described toxic BXD aglucone DIMBOA. The better-performing corn strain had higher re-glucosylation activity towards DIMBOA, catalyzed by SfUGT33F28, than the poorer-performing rice strain. The two strains also exhibit differential UGT regulation at the transcript level in response to DIMBOA. Our previous work with FAW had demonstrated that the gene encoding the principal DIMBOA re-glucosylating UGT, *UGT33F28*, showed a constitutively higher expression of the canonical transcript in the corn strain over the rice strain (accompanied by higher specific activities towards DIMBOA) and an up-regulation of the transcript in both the strains upon a diet shift from a pinto bean based artificial diet to a maize leaf diet (Israni et al. 2020). Subsequent work has provided evidence for the expression of UGT33F28 splicing variants that can regulate the reglucosylation activity of the canonical protein. The expression of transcript variant 1 (*tv1*) in particular was demonstrated to be induced in response to DIMBOA treatment, with the translated protein SfUGT33F28_v1 physically interacting with, and positively regulating the activity of the canonical protein (Israni et al. 2022). Transcript variant 1 expression was constitutively higher in rice strain than corn strain larvae. On the population level, rice strain population RS-1 showed higher expression compared to population RS-2, while the canonical transcript was comparable in both populations. Accordingly, population RS-1 larvae exhibited a much higher glucosylation activity towards DIMBOA compared to population RS-2 larvae. Eventually, the ratio of *UGT33F28* variants remains critical, so a much higher constitutive as well as induced expression of the canonical transcript in the corn strain larvae explains its greater enzymatic activity towards DIMBOA compared to the rice strain. Expression of these variants could nevertheless enable the rice strain larvae to enhance their overall detoxification capability upon maize feeding.

Detoxification of plant defenses could play a major role in FAW specialization on other host plants besides maize. For example, in FAW feeding on rice, members of the cytochrome P450 family *SfCYP321A9* and *SfCYP9A58* have been shown to mitigate the effects of ferulic acid (a hydroxycinnamic acid), gramine (an indole alkaloid) and tricin (an O-methylated flavone), since their knockdown leads to a prolonged development for FAW larvae on rice plants (He et al. 2023). In another study, knockdown of *SfCYP12A2* and *SfUGT41B8* had important consequences for larval adaptation on rice. Both the genes were differentially expressed in the two FAW strains and *SfCYP12A2* knockdown in particular led to a decrease in toleration of rice flavonoid luteolin (Han et al. 2023).

Other factors could influence the specialization of FAW strains on their host plants, such as the gut microbiome. FAW strains sampled from across the Western Hemisphere show no clear differences due to strain or geographical distribution of the populations analyzed (Oliveira, Rodrigues,Cônsoli 2023). On the other hand, another recent study showed that switching the FAW strains kept on artificial diet to either corn or rice as host and maintaining them for several generations led to the dominance of corn strain gut by Firmicutes and a contrasting dominance of *Enterococcus* in the rice strain gut (Han et al. 2023).

Taken together, we have identified several factors that might account for the better performance of the corn FAW strain on its characteristic host plant over the rice strain. Reduction of plant defense signaling, reduced BXD accumulation and a greater BXD detoxification potential could all add up to improving insect performance. Additional factors, such as variation observed in the accumulation of other defense metabolites, and a more appropriate gut microbiome should be explored in future studies.

## Methods

### Insects and plants

Two host strains of FAW were used in the present study with two populations each: corn strain 1 (PRC; CS-1) and 2 (COBS; CS-2) and rice strain1 (FLR; RS-1) and 2 (ONA; RS-2). CS-1 was originally sourced from maize in Puerto Rico (Santa Isabella) in 2010, CS-2 from maize in Florida (Citra) in 2018. RS-1 was collected in pasture in Florida (Moore Haven) in 2010, RS-2 from grassland in Florida (Ona) in 2019. All strains were established as lab strains at the Department of Entomology at the Max Planck Institute for Chemical Ecology (MPI-CE). Collected larvae were genotyped using strain-specific cytochrome oxidase I gene (COI) markers (all strains) (Hänniger et al. 2017) and additionally by strain-specific Triosephosphate isomerase (Tpi) SNPs (CS-2 and RS-2). Lab colonies were maintained with single pair matings, reared on a semi-artificial diet based on pinto bean, and under controlled light and temperature conditions (16:8 h light/dark, 26 °C).

Seeds of *Z. mays* were obtained commercially (Badischer Gelber variety, Kiepenkerl, Germany), and grown under controlled light and temperature conditions (16:8 h light/dark, daytime temperature 22 °C, nighttime 20 °C).

### Feeding experiments

Second or third instar FAW larvae were utilized for all experiments. Insects were starved overnight prior to feeding experiments. The following day insects were put on maize plants secured with perforated plastic bags. The larvae were allowed to feed for a duration of 1-24 hours. Each plant was treated with one caterpillar in a completely randomized design. Leaves were collected to account for both local and systemic effects of FAW herbivory. Leaf material from different treatment groups was harvested and flash-frozen in liquid nitrogen. Insects that had fed on maize leaves or artificial diet were used for isolation of guts using cold phosphate buffered saline (pH 7.4), and the guts were subsequently cleaned and flash-frozen in liquid nitrogen until further use.

### Maize leaf sample preparation

For measurements of maize metabolites, including untargeted measurements and targeted measurements of BXDs, extracts were prepared using leaf samples obtained from plants used from the feeding experiments. Samples were ground using liquid nitrogen, quickly transferred to tubes, freeze dried, weighed and extracted using freshly prepared acidified water/methanol (1:1, v:v, pH 3 using 0.5% formic acid). Extracts were vortexed for 2 min, agitated in a paint shaker for 2 min (with 3 mm steel beads), centrifuged at 16,000 g for 5 minutes, and the obtained supernatants were collected and analyzed by LC-MS/MS. For targeted phytohormone measurements, these methanolic extracts were spiked with an internal standard mix comprising 40 ng D6-JA, 8 ng D6-JA-Ile (both HPC Standards GmbH, Germany), 40 ng D4-SA (Santa Cruz Biotechnology, USA), and 40 ng D6-ABA (Toronto Research Chemicals, Canada); and subsequently analyzed by LC-MS/MS.

### RNA extraction, reverse transcription and real time-PCR analysis

Maize tissue samples homogenized and total RNA extracted using the Qiagen RNeasy Plant Mini kit. RNA concentrations were measured with the NanoDrop 2000 UV-Vis Spectrophotometer (Thermo Scientific). Insect RNA preparation was carried out using innuPREP RNA Mini Kit (Analytik Jena). First strand cDNA was synthesized from 1 μg total RNA using SuperScript III Reverse Transcriptase and OligodT primers from Invitrogen. Real time PCR analyses were carried out using Brilliant III SYBR Master Mix (Agilent), employing SYBR Green chemistry. Relative quantification of the transcript levels was done using the 2-ΔΔCt method. *Z. mays actin* was used as reference gene for all analyses performed with maize plants, and *S. frugiperda RPL10* used as a reference gene for all analyses performed with FAW larvae. The primer pairs used for analyzing the transcripts are listed in Supplementary Table 1 (Ahmad et al. 2011; Handrick et al. 2016; Israni et al. 2022; Sun et al. 2021)). The specificity of the primer pairs was confirmed by sequencing. The real time PCR products were electrophoresed on a 2% agarose gel, followed by clean up using the PCR clean up kit (Qiagen), cloning into pCR-Blunt II-TOPO vector (Life Technologies) and transformation into NEB ready to use competent cells (Life Technologies), which were plated on selective LB agar medium containing 100 μg/mL ampicillin and incubated overnight at 37 °C. Colonies were cultured overnight for plasmid isolation and the plasmids were confirmed by sequencing.

### Enzymatic assays

For conducting UDP-glucosyltransferase (UGT) assays from the different FAW strains, caterpillars were dissected in cold phosphate buffered saline (pH 7.4), and their gut tissues were collected, cleaned of any diet and homogenized. A typical enzyme reaction included 10 μL homogenate, 2 μL of 12.5 mM DIMBOA in DMSO (25 nmol), 4 μL of 12.5 mM UDP-glucose in water (50 nmol), and phosphate buffer (pH 7.0, 100 mM) to give an assay volume of 50 μL. Controls containing either boiled enzymatic preparation, or those containing only the sample and buffer were included. After incubation at 30°C for 60 min, the enzyme reactions were interrupted by adding 50 μL of a methanol: formic acid 1:1 (v:v) solution. Assay tubes were centrifuged at 5,000 *g* for 5 min and the obtained supernatant was collected and analyzed by LC-MS/MS.

### Chemical analysis

For the untargeted analyses, ultra-high-performance liquid chromatography– electrospray ionization– high resolution mass spectrometry (UHPLC–ESI–HRMS) was performed with a Dionex Ultimate 3000 series UHPLC (Thermo Scientific) and a Bruker timsToF mass spectrometer (Bruker Daltonics, Germany). UHPLC was used applying a reversed-phase Zorbax Eclipse XDB-C18 column (100 mm × 2.1 mm, 1.8 μm, Agilent Technologies, Germany) with a solvent system of 0.1% formic acid and acetonitrile as mobile phases A and B respectively at a flow rate of 0.3 mL/min. The elution profile was the following: 0 to 0.5 min, 5% B; 0.5 to 11.0 min, 5% to 60% B in A; 11.0 to 11.1 min, 60% to 100% B, 11.1 to 12.0 min, 100% B and 12.1 to 15.0 min 5% B. Electrospray ionization (ESI) in negative or positive ionization mode was used for the coupling of LC to MS. The mass spectrometer parameters were set as follows: capillary voltage 4.5 KV/3.5KV, end plate offset of 500V, nebulizer pressure 2.8 bar, nitrogen at 280°C at a flow rate of 8L/min as drying gas. Acquisition was achieved at 12 Hz with a mass range from *m*/*z* 50 to 1500, with data-dependent MS/MS and an active exclusion window of 0.1 min, a reconsideration threshold of 1.8-fold change, and an exclusion after 5 spectra. Fragmentation was triggered by an absolute threshold of 50 counts and acquired on the two most intense peaks with MS/MS spectra acquisition of 12 Hz. Collision energy was alternated between 20 and 50V. At the beginning of each chromatographic analysis, 10 μL of a sodium formate-isopropanol solution (10 mM solution of NaOH in 50/50 (*v*/*v*%) isopropanol water containing 0.2% formic acid) was injected into the dead volume of the sample injection for re-calibration of the mass spectrometer using the expected cluster ion *m*/*z* values. Peak detection was carried out using Metaboscape software (Bruker Daltonik, Germany) with the T-Rex 3D algorithm for qTOF data. For peak detection the following parameters were used: intensity threshold of 700 with a minimum of 10 spectra, time window from 0.4 to 11.8 min, peaks kept if they were detected in at least all replicates of one sample group.

For all targeted analytical procedures, formic acid (0.05%) in water and acetonitrile were used as mobile phases A and B, respectively, and the column temperature was maintained at 25 °C. Analyses of plant samples used a Zorbax Eclipse XDB-C18 column (50 x 4.6 mm, 1.8 μm, Agilent Technologies) with a flow rate of 1.1 mL/min and with the following elution profile: 0-0.5 min, 95% A; 0.5-6 min, 95-67.5% A; 6.02-7 min, 100% B; 7.1-9.5 min, 95% A. For BXD quantification, LC-MS/MS analyses were performed on an Agilent 1200 HPLC system (Agilent Technologies) coupled to an API 6500 tandem spectrometer (AB Sciex) equipped with a turbospray ion source operating in negative ionization mode. Multiple reaction monitoring (MRM; see Supplementary Table 2) was used to monitor analyte parent ion to product ion conversion with parameters from the literature (Israni et al. 2020; Pedersen et al. 2011; Wouters et al. 2014). BXD standards [(2*R*)-DIMBOA-Glc and MBOA-Glc] were kindly provided by Prof. Dieter Sicker (University of Leipzig, Germany), and Dr. Gaétan Glauser (University of Neuchâtel, Switzerland) respectively.

Phytohormone analysis was performed on an Agilent 1200 series HPLC system (Agilent Technologies) as described previously (Heyer, Reichelt, Mithöfer 2018), with the modification that a tandem mass spectrometer QTRAP 6500 (SCIEX, Darmstadt, Germany) was used in multiple reaction monitoring (MRM) mode with parameters listed in Supplementary Table 3. Concentrations of *cis*-OPDA, and OH-JA were determined relative to the quantity of the internal standard D6-JA applying a response factor (RF) of 1.0, while OH-JA-Ile and COOH-JA-Ile were quantified relative to D6-JA-Ile, applying a response factor (RF) of 1.0. Since we observed that both the D6-labeled JA and D6-labeled JA-Ile standards (HPC Standards GmbH, Germany) contained 40 % of the corresponding D5-labeled compounds, the sum of the peak areas of D5- and D6-compound was used for quantification. Analyst (version 1.6.3, Applied Biosystems) software was used for data acquisition and processing.

### Statistical analysis

MetaboAnalyst 5.0 was used to perform all the analyses for untargeted metabolomics and generate figures for data visualization (Chong, Wishart, Xia 2019). Data were suitably transformed, and scaled using Pareto scaling. False discovery rate was applied to adjust the *p*-values (0.05). Pairwise analysis was performed using volcano plots, which combine fold change (FC ≥ |2.0|) and t-test analysis, which can control the false discovery rate (p ≤ 0.05), to identify the features that were potentially significant in discriminating the strains on maize plants. All other statistical analyses on targeted metabolite measurements and gene expression were carried out using SigmaPlot 12.0 and R studio (version 3.6.3). Data were tested for homogeneity of variance and normality and were appropriately transformed to meet these criteria where required to carry out group comparisons. The specific statistical method used for each data set is described in the figure legends. All graphs were created using R studio.

## Acknowledgments.

The authors would like to thank Melanie Unbehend, Rob Meagher and Jehangir Bhadha for their help with collecting the PRC and FLR (MU), COBS (RM) and ONA (JB) strains and André Richter and Angelika Berg for assistance with insect rearing; the greenhouse team of the Max Planck Institute for Chemical Ecology for growing maize plants and Felipe C. Wouters for earlier work generating methods utilized in this project. The DFG projects HA 8327/1-1 and HA 8327/2-1 provided funding for insect collection, genotyping and maintenance. The Max Planck Society provided funding for publication and open access via Project DEAL.

## Author contributions

B.I. and D.G.V. conceived and designed the experiments. S.H. field collected, genotyped and maintained insects. B.I. performed FAW herbivory experiments, processed samples for metabolomics, for RNA isolation and cDNA preparation; B.I. and M.R. conducted metabolomics analyses and data processing; B.I. and B.R. performed real time PCR analyses; B.R. performed sequencing to confirm real time PCR primer specificity. B.I., J.G. and D.G.V. wrote the manuscript with input and approval from all authors.

## Supplementary data

**Supplementary figure 1.**
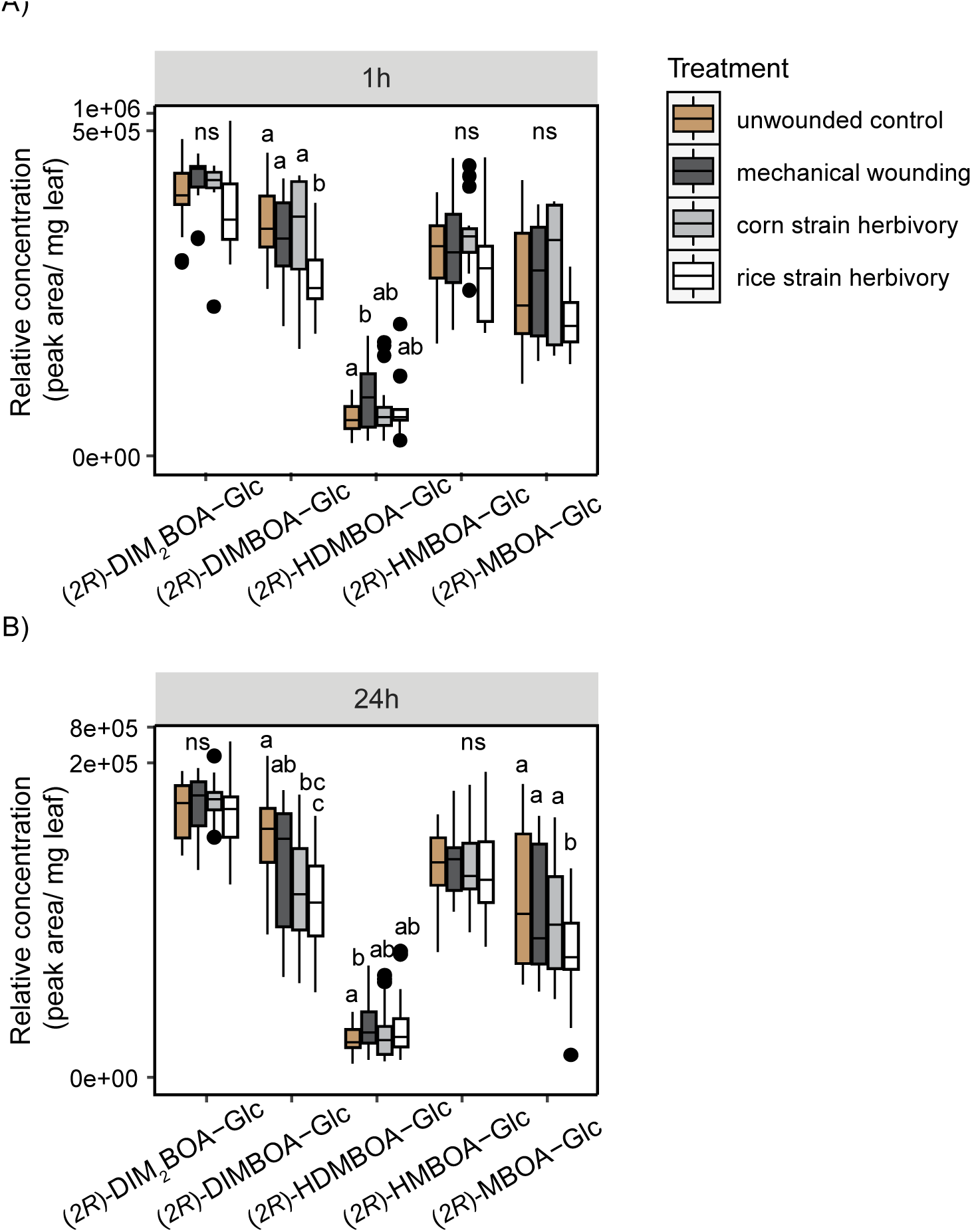
BXD biosynthetic pathway activation in maize plants in response to FAW herbivory. Relative BXD glucoside accumulation in maize leaves subject to FAW herbivory expressed as relative concentration (peak area/mg leaf) (n= 30-40). Two-way ANOVA was performed and Holm Sidak method was applied to perform pairwise comparisons. All data were log transformed to meet the criteria for normality for conducting statistical tests. Small letters on the plots represent significant differences (P< 0.05). (*2R*)-DIM_2_BOA-Glc, 2-(2,4-dihydroxy-7,8-dimethoxy-1,4-benzoxazin-3-one)-β-d-glucopyranose; (*2R*)-DIMBOA-Glc, 2-(2,4-dihydroxy-7-methoxy-1,4-benzoxazin-3-one)-β-d-glucopyranose; (*2R*)-HDMBOA-Glc, 2-(2-hydroxy-4,7-dimethoxy-1,4-benzoxazin-3-one)-β-d-glucopyranose ; (*2R*)-HMBOA-Glc, 2-(2-hydroxy-7-methoxy-1,4-benzoxazin-3-one)-β-d-glucopyranose; and (*2R*)-MBOA, 2-(6-methoxy-2-benzoxazolinone)-β-d-glucopyranose.

**Supplementary Table 1.**
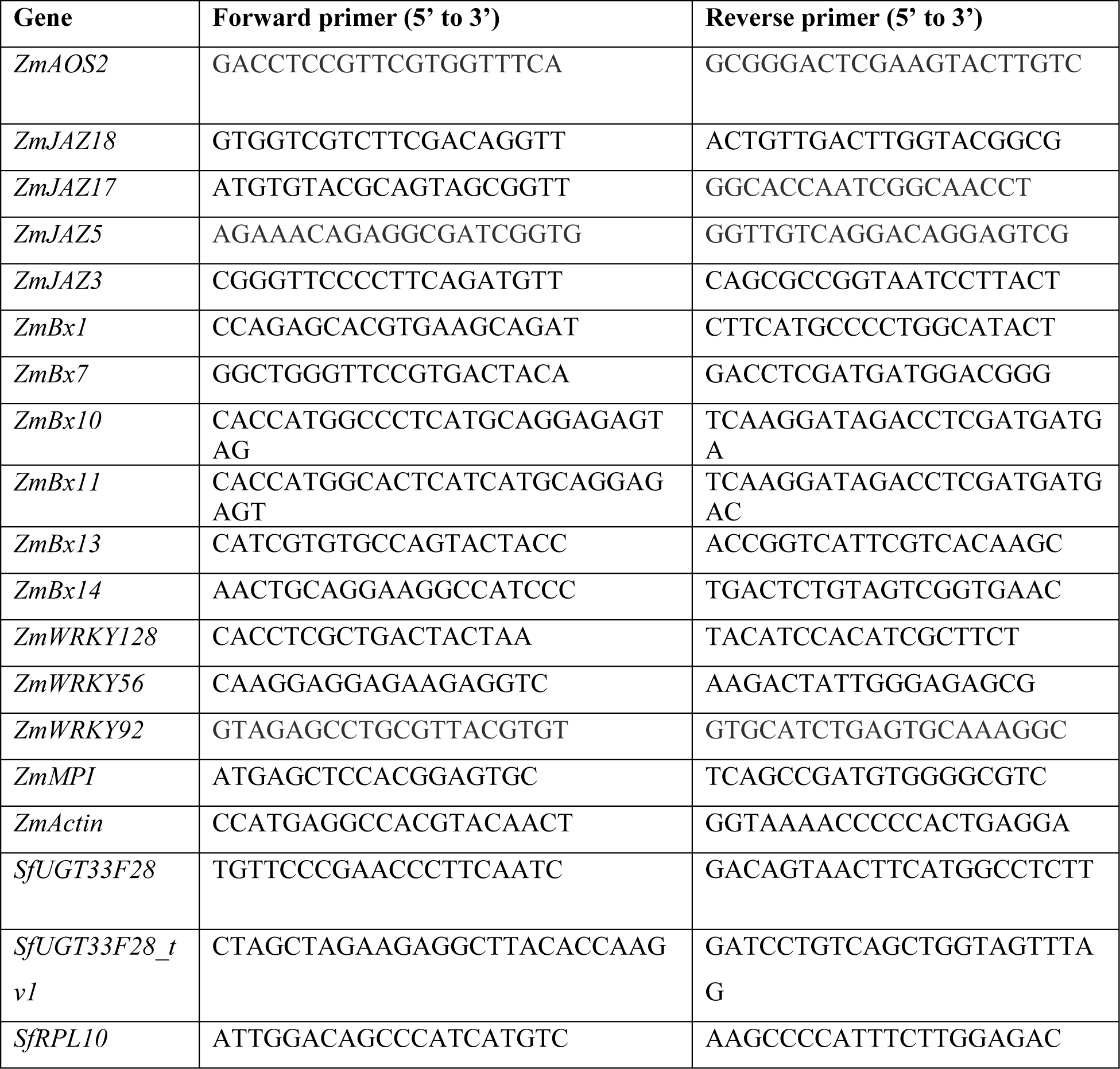
List of primers utilized for real time PCR analyses.

**Supplementary Table 2.** Details of the analysis of Benzoxazinoids (BXDs) by LC-MS/MS. BXDs were measured using an Agilent HPLC 1200/API6500 (Sciex) instrument in negative ionisation mode. Abbreviations are: Q1, selected *m/z* of the first quadrupole; Q3, selected *m/z* of the third quadrupole; DP, declustering potential (V); and CE, collision energy (V).

| Q1 | Q3 | Dwell<br>time (sec) | Compound | DP | CE |
| --- | --- | --- | --- | --- | --- |
| 418 | 372 | 10 | DIMBOA-Glc | -40 | -18 |
| 210 | 149 | 10 | DIMBOA | -40 | -16 |
| 342 | 134 | 10 | DIBOA-Glc | -40 | -24 |
| 180 | 134 | 10 | DIBOA | -40 | -10 |
| 448 | 402 | 10 | DIM<sub>2</sub>BOA-Glc | -40 | -18 |
| 240 | 179 | 10 | DIM<sub>2</sub>BOA | -40 | -16 |
| 326 | 164 | 10 | HBOA-Glc | -40 | -20 |
| 164 | 108 | 10 | HBOA | -40 | -23 |
| 194 | 138 | 10 | HMBOA | -40 | -19 |
| 432 | 356 | 10 | HDMBOA-Glc | -40 | -20 |
| 224 | 164 | 10 | HDMBOA | -40 | -15 |
| 462 | 194 | 10 | HDM<sub>2</sub>BOA-Glc | -40 | -25 |
| 254 | 179 | 10 | HDM<sub>2</sub>BOA | -40 | -15 |
| 164 | 149 | 10 | MBOA | -40 | -20 |
| 356 | 194 | 10 | HMBOA-Glc | -40 | -15 |
| 194 | 164 | 10 | M<sub>2</sub>BOA | -40 | -20 |
| 402 | 194 | 10 | DIM<sub>2</sub>BOA -Glc II | -40 | -18 |
| 372 | 164 | 10 | MBOA-Glc II | -40 | -20 |
| 372 | 326 | 10 | MBOA-Glc | -40 | -12 |
| 372 | 210 | 10 | DIMBOA-Glc II | -40 | -15 |
| 402 | 356 | 10 | HMBOA-Glc II | -40 | -18 |

**Supplementary Table 3.** Details of the analysis of phytohormones by LC-MS/MS using an Agilent HPLC 1260/QTRAP6500 (Sciex) instrument in negative ionisation mode. Abbreviations are: Q1, selected *m/z* of the first quadrupole; Q3, selected *m/z* of the third quadrupole; RT, retention time; DP, declustering potential (V); and CE, collision energy (V); SA, salicylic acid; JA, jasmonic acid; ABA, abscisic acid; JA-Ile, jasmonic acid-isoleucin conjugate; OPDA, 12-oxo phytodienoic acid; OH-JA, 12-hydroxy-jasmonic acid; OH-JA-Ile, 11/12-hydroxy-jasmonic acid-isoleucin conjugate; COOH-JA-Ile, 11/12-carboxy-jasmonic acid-isoleucin conjugate; D4-SA, D4-salicylic acid; D6-JA, D6-jasmonic acid; D5-JA, D5-jasmonic acid; D6-ABA, D6-abscisic acid; D6-JA-Ile, D6-jasmonic acid-isoleucin conjugate; D5-JA-Ile, D5-jasmonic acid-isoleucin conjugate.

| Q1 | Q3 | RT (min) | Compound | Internal standard | DP | CE |
| --- | --- | --- | --- | --- | --- | --- |
| 136.93 | 93 | 3.3 | SA | D4-SA | -20 | -24 |
| 209.07 | 59 | 3.6 | JA | D6-JA | -20 | -24 |
| 263 | 153.2 | 3.4 | ABA | D6-ABA | -20 | -22 |
| 322.19 | 130.1 | 3.9 | JA-Ile | D6-JA-Ile | -20 | -30 |
| 290.9 | 165.1 | 4.6 | OPDA | D6-JA | -20 | -24 |
| 225.1 | 59 | 2.6 | OH-JA | D6-JA | -20 | -24 |
| 338.1 | 130.1 | 3.0 | OH-JA-Ile | D6-JA-Ile | -20 | -30 |
| 352.1 | 130.1 | 3.0 | COOH-JA-Ile | D6-JA-Ile | -20 | -30 |
| 140.93 | 97 | 3.3 | D4-SA |  | -20 | -24 |
| 215 | 59 | 3.6 | D6-JA |  | -20 | -24 |
| 214 | 59 | 3.6 | D5-JA |  | -20 | -24 |
| 269 | 159.2 | 3.4 | D6-ABA |  | -20 | -22 |
| 328.19 | 130.1 | 3.9 | D6-JA-Ile |  | -20 | -30 |
| 327.19 | 130.1 | 3.9 | D5-JA-Ile |  | -20 | -30 |

